# Reciprocal interactions between the sorghum root microbiome and the parasitic weed *Striga hermonthica*

**DOI:** 10.1101/2023.10.01.560362

**Authors:** Desalegn W. Etalo, Dominika Rybka, Lorenzo Lombard, Marcio F.A. Leite, Adam Ossowicki, Francisco Dini-Andreote, Luisa Arias-Giraldo, Eiko E. Kuramae, Pedro W. Crous, Taye Tessema, Jos M. Raaijmakers

## Abstract

The soil microbiome plays a crucial role in protecting plants against various pests and pathogens. However, its impact on interactions between plants and parasitic weeds, such as *Striga hermonthica*, is poorly understood. In this study, sorghum plants susceptible to *Striga* were grown in 22 different field soils infested with parasite seeds. Significant variations in *Striga* infections were observed among the soils. When the most *Striga*-suppressive soil was gamma-irradiated, there was a significant increase in *Striga* attachments, highlighting the importance of the soil microbiome in disrupting parasite infection. In the presence of the soil microbiome, the *Striga*-susceptible sorghum plants performed similarly to three *Striga*-resistant genotypes. This effect was lost when the soil microbiome was eliminated by gamma-irradiation. Subsequent analysis revealed that *Striga* substantially affected the sorghum rhizosphere microbiome and that both the sorghum rhizosphere mycobiome and bacteriome composition significantly correlated with *Striga* attachment. Interestingly, certain fungal species in the sorghum rhizosphere mycobiome were only detected when *Striga* seeds were present. Further investigation showed that these fungal taxa originated from the *Striga* seeds and are known sorghum pathogens, suggesting a potential partnership between *Striga* and fungal pathogens to invade their shared host. Overall, our study demonstrated that the soil microbiome influences *Striga* infection and sorghum performance in a genotype-dependent manner, and the microbiome of *Striga* seeds affects the composition of the sorghum rhizosphere microbiome.

## Introduction

The hemiparasitic weed *Striga hermonthica* has long impacted diverse crop production systems in sub-Saharan Africa. Over 60% of the cultivated farmlands in this region are infested with one or more species of *Striga*, causing substantial yield losses up to complete crop failure (*1*). Moreover, *Striga* is expanding its host range and is steadily increasing its geographical distribution (*1*). Maize, sorghum, millet, and rice are the most important cereal crops in sub-Saharan Africa affected by *Striga.* Beyond its impact on food security, *Striga* is of significant biodiversity and ecological concern as it leaves the farmers in sub-Saharan Africa no choice but to abandon their farms and encroach into uncultivated lands and natural reserves.

Compared to the self-crossing *Striga asiatica*, the obligate outbreeding *Striga hermonthica* exhibits a higher within-population genetic diversity (*2*, *3*). In addition, *S. hermonthica* produces 10,000-85,000 seeds per emerged plant that remain viable in the soil for several years (*4–6*). Under favorable environmental conditions and host-derived chemical stimuli (i.e., strigolactones), *Striga* seeds germinate, produce a haustorium, attach to the host roots, establish a connection with the vascular host tissue, and then siphon water, essential nutrients, and photo-assimilates, thereby debilitating crop growth and yield. Most damage to the host occurs during the early stages of parasitism when *Striga* is in the subterranean stages of its life cycle (*7*). The high seed production, the longevity of the *Striga* seed bank, its vast genetic diversity, and the intimate association of the *Striga* life cycle with the host physiology are significant constraints in developing effective strategies to control this parasite. Hence, intervention strategies that deplete the seed bank and disrupt the early stages of infection are preferred effective control strategies.

Multiple intervention strategies have been pursued, including using non-host plants and synthetic chemicals (e.g., ethylene, strigolactone analogs) to induce suicidal seed germination without the host (*8–12*). Another viable approach to disrupt the early stages of infection is breeding for *Striga* resistance in the host (*13*). To date, host genetic traits for low *Striga* germination stimulant (*LGS*) production, low production of haustoria initiation factors (*LHF*), hypersensitive response (HR), and incompatible response (IR) to parasite invasion are the main targets of resistance breeding (*13*). However, the major challenges for sorghum breeding programs are inconsistent durability and stability of host resistance in different agroecological zones and against genetically diverse *Striga* ecotypes (*14–18*). Hence, the selection of durable resistance for *Striga* should consider the host genotype, the parasite genotype, and the environment. For the environmental component, most studies have focused on soil chemical properties, particularly phosphorus and nitrogen (*19*–*21*), with little to no attention to the soil microbiome.

Owing to their importance in protection against pathogens, insects, and nematodes (*22–25*), soil and plant microbiomes are essential for the sustainable management of biotic and abiotic stresses. In the past decades, however, most studies on host-parasite interactions focused only on host genetics, biochemistry, and physiology, keeping the microbiome aside as a source of unmeasured host genetic variation. Hence, the soil and root microbiome’s contribution to plant-parasitic weed interactions remains largely unexplored (*26*). The interaction between the host and *Striga* mainly occurs in the rhizosphere, where many microbes reside and are active. We hypothesized that in the co-evolutionary arms race between plants and parasitic weeds, both parties could have developed the ability to modulate the microbiome to strengthen either host resistance or parasite invasion. Here, we report for the first time the impact of the soil microbiome on *Striga* infection in both *Striga*-susceptible and resistant sorghum genotypes. Furthermore, we show that *Striga* infection and the microbiome of *Striga* seeds reciprocally affect the sorghum root microbiome composition. The potential implication of these results for future *Striga* intervention strategies is discussed.

## Materials and Methods

### Soil collection, history, and physicochemical properties

Twenty-two predominantly agricultural fields with diverse cropping histories were sampled in the Netherlands. We chose field soils in the Netherlands instead of soils from sub-Saharan Africa to avoid the confounding effects of the indigenous *Striga* seed bank in many African sorghum field soils. The cropping history and physicochemical properties of each field soil are provided in **Supplementary Table ST1** and **Supplementary Figure SF1**. The soils were air-dried, sieved through a 4-mm mesh to remove stones and plant debris, and stored at 8 °C until further use.

### *Striga* seed collection and grading

*Striga hermonthica* seeds were collected from infested sorghum fields in the Abergele zone of the Tigray regional state of Ethiopia. Seeds were cleaned from debris using multilayer sieves (420, 180, and 150 µm), and *Striga* seeds captured on the 180 µm sieve were used for this study. The purity, germination percentage, and the number of *Striga* seeds per mg were determined following standard procedures and used to determine the number of germinable seeds required for the *Striga* infection assay (*27*).

### Seed surface sterilization

Sorghum seeds (cultivars: Birhan (PSL5061), Framida, Gubiye (P9401), SRN-39, and Shan Quei Red (SQR)) were washed with sterile water and then surface sterilized for 20 min with 1.5% (v/v) NaOCl solution under continuous shaking. After discarding the solution, the seeds were washed three times with sterile water. *Striga* seeds were initially washed with sterile water and surface sterilized for 4 min with 1.0% (v/v) NaOCl solution supplemented with 0.01% (v/v) Tween 80 under continuous shaking. After discarding the NaOCl solution, the seeds were washed thrice with sterile water. For both sorghum and *Striga* seeds, 100 µL of the last washes were plated on MEA and 1/10^th^ TSA agar plates to assess the effectiveness of the surface sterilization.

### Preconditioning of *Striga* seeds and pre-germination of sorghum seeds

The surface-sterilized seeds were mixed thoroughly with 1 kg of non-sterile fine sand in an opaque container, and the moisture content was adjusted to 10% (v/w) with tap water and incubated at 28 °C for 14 days. For pre-germination of the surface-sterilized sorghum seeds, double-layered 85-mm-diameter sterile glass fiber filter papers were aligned in a 90-mm-diameter sterile Petri dish, and 5 mL of sterile water was added. The sorghum seeds were placed on the wet glass fiber filter paper, and the plate was sealed with Parafilm and incubated at 28 °C for 24 h.

### Bioassay to investigate sorghum-*Striga*-microbiome interactions

The soil plug system developed in this study is composed of three elements: i) a plug of the field soil, ii) river sand as a standardized background substrate with or without pre-conditioned *Striga* seeds, and iii) the host plant sorghum (**Supplementary Figure SF2a**). The soil plug represents 1% (v/v) of the background substrate and serves as the source of indigenous soil microbes colonizing the roots of the sorghum seedlings. The moisture content of the soils was raised to 10% and incubated for two days at 28 °C. The sorghum roots grow through the field soil plug into the background substrate amended with or without (control) *Striga* seeds. When screening multiple field soils, the soil plug system reduces the confounding influence soil physicochemical properties may have on plant growth and root development. Furthermore, the river sand as the background substrate amended with (or without) *Striga* seeds made it feasible to remove from the sorghum roots, thereby facilitating the counting of the root infections caused by *Striga*.

For the screening of the 22 field soils, we changed the background medium to a mixture of fine- and medium-sized sand (1:1 w/w) with the addition of 5,000 germinable preconditioned *Striga* seeds per pot for the *Striga* infection bioassays. The sand and the conditioned *Striga* seeds were homogenized in a concrete mixer by adjusting the moisture content of the medium to 8% (v/w) with tap water. The pots were filled with the *Striga* seed-sand mixture, and a hole of 3 mm diameter and 6 cm depth was made for the soil plug consisting of 30 g of the respective soils with a moisture content of 10% (v/w). One pre-germinated sorghum seed was planted in the center of each soil plug. The plants were grown at 25 °C in the greenhouse with a 12/12 h day/night cycle for six weeks. Supplemental lighting was provided when needed during the growing period. The relative air humidity was set at 60%.

Half-strength Hoagland solution with reduced phosphate (10% of the recommended dose) was provided to the plants two and four weeks after planting. Watering of the plants was done every two days and was stopped seven days before harvesting. For the harvest, plants were carefully removed from the pots in a tray lined with tissue paper, the roots were gently separated from the sand, and the number of *Striga* attachments per plant was scored. Next, the shoot was cut at the collar, and the roots were shaken to remove the loosely adhering sand particles and transferred to a sterile glass container with 100 mL of sterile water. The bottle was closed and kept at 4 °C until further processing. Three biological replicates were considered for each soil sample.

### Root colonization proof of principle assay

To track the longitudinal colonization of sorghum root by microbes from a soil plug, we introduced GFP-labeled and rifampicin-resistant rhizobacteria *Pseudomonas fluorescens* strain SS101 into soil plugs at a concentration of 10^6^ cfu/g of soil in natural and gamma-irradiated soil. Surface sterilized and pregerminated sorghum seeds were planted in the soil plugs and grown under standard conditions in the greenhouse for two weeks. Roos were sampled at 10, 20, and 30 cm (**Supplementary Figure SF2b**) from the plug and kept in 0.9 % NaCl solution. The root samples were shaken at 250 rpm for 30 min, and the suspension was plated at different dilutions on LA medium supplemented with rifampicin. The number of colony-forming units was counted.

### Rhizosphere soil collection and DNA extraction

The roots were kept in closed glass containers with 100 mL water and shaken at 250 rpm for 1 hr, sonicated for 10 min, and shaken at 250 rpm for 8 min. For making glycerol stocks, 10 mL of the solution was transferred to a 15 mL falcon tube and centrifuged at maximum speed for 10 min. Next, the supernatant was removed, the pellet was re-suspended in 3 mL 10 mM MgSO_4_ solution, and an equal portion of the suspension was mixed with sterile 80% (v/v) glycerol to constitute a final concentration of 40% (v/v). Finally, the remaining rhizosphere suspension was divided into two 50 mL Falcon tubes and freeze-dried for total DNA extraction.

DNA was extracted from the freeze-dried samples using a DNeasy® PowerSoil® Pro Kit according to the manufacturer’s guidelines with slight modifications from the manufacturer (*28*). Briefly, 600 µL of the lysis buffer was added to the freeze-dried rhizosphere soil in a 50 mL Falcon tube and vortexed for 30 sec. Then, the mixture was transferred to the power bead tube and subjected to the Powerlyzer for 10 min at max speed. After eluting the DNA with 50 µL C6 buffer, its concentration and quality were determined by Nanodrop.

### *Striga* seed microbiome analyses

To analyze the *Striga* seed bacterial and fungal communities, 400 mg of *Striga* seeds were washed in sterile water. After discarding the water, the seeds were resuspended in 0.85% (w/v) NaCl solution and shaken for 3 hrs at 100 rpm. Next, the suspension was transferred to a new falcon tube, freeze-dried, and kept at -20 °C until further use. To extract the endophytic microbes (i.e., inside the *Striga* seeds), the seeds were first washed with sterile water and surface sterilized with 1% (v/v) NaOCl solution for 4 min. This was followed by treatment with 70% (v/v) ethanol and washed thrice with sterile water. Then, the seeds were suspended in 0.85% (w/v) NaCl and crushed under sterile conditions with a mortar and pestle. Next, the crushed seeds were transferred to a new falcon tube, sonicated for 10 min, and vortexed for 15 min. The sample was then filtered through 75 μm sterile mesh to remove large seed fragments, and the filtrate was freeze-dried and kept at -20 °C. Finally, as stated above, the freeze-dried samples containing the *Striga* seed microbiome’s epiphytic and endophytic fractions were subjected to DNA extraction. Three replicate seed samples were considered for the epiphytic and endophytic seed microbiome analyses.

### Amplicon sequencing of the sorghum rhizosphere and *Striga* seed microbiomes

The bacterial and fungal communities of the sorghum rhizosphere were characterized by sequencing amplicons of the V3-V4 region of 16S rRNA and the ITS2 region of the Internal Transcribed Spacer (ITS) genes, respectively, using Illumina MiSeq at BaseClear (Leiden, Netherlands). While for the Striga seeds, the V3-V4 amplicon of the 16S rRNA and the ITS1 amplicon regions were sequenced using Illumina MiSeq at Genome Quebec (Canada). Sequencing primers were removed with Cutadapt (*29*) from the FASTQ files. The 16S rRNA gene and the ITS sequences were processed using DADA2 (*30*) in the Qiime2 platform (*31*), and an amplicon sequence variant (ASV) table was generated. The ASVs were further clustered to OTUs with 97% similarities. The full-length Silva 138.1 reference sequences were used to generate a classifier file for the V3-V4 regions of the 16S rRNA, and the UNITE (ver9_97_s_29.11.2022) databases were used to generate a classifier file for ITS sequences. The two classifier files were used for the taxonomic assignment of the Operational Taxonomic Units (OTUs) for 16S rRNA and ITS1/2 sequences. The link for the corresponding code for the data analysis is provided in the data availability section of the manuscript.

### Characterization of fungi from the sorghum rhizosphere and *Striga* seeds

Different dilutions of the sorghum rhizosphere soil stored in glycerol stocks were plated on MEA-medium supplemented with chloramphenicol (25 µg/mL). For the fungal isolation from *Striga* seeds, seeds were surface-sterilized to isolate endophytic fungi. Individual seeds were crushed with a tweezer and plated on MEA medium amended with streptomycin and penicillin (100 µg/mL) to suppress bacterial growth. The fungal cultures were purified by hyphal tip isolation. According to the manufacturer’s instructions, DNA was isolated from 7-day-old axenic fungal cultures grown on MEA using the Wizard® Genomic DNA purification Kit (Promega Corporation, Madison, WI, USA). The sequences of the following gene regions were determined using the primer sets and conditions as described in (*32*): both the Internal Transcribed Spacer regions and intervening 5.8S (ITS), 28S nrDNA (LSU), translation elongation factor–1α (*TEF*–1α), DNA-directed RNA polymerase II largest (RPB1 and RPB2) β-tubulin (*tub2*) and Calmodulin (CAL) gene regions. All amplicons generated were sequenced in both directions with the same primer sets as for amplicon analyses. Consensus sequences for each locus were determined using MEGA v. 7 (*33*) and megablasted against the NCBI’s GenBank nucleotide database. Taxonomic delineation of the isolated fungi was further confirmed using the Westerdijk Institute in-house database for fungal identification. The sequence data obtained for these markers are available in **Supplementary Table ST9**, which can be found at the following link: [https://github.com/DesalegnE/Reciprocal-interactions_Etalo-et-al-2023-].

### Statistical analysis

Univariate analysis

#### Screening of field soils for Striga suppression

The influence of soil samples (independent variables) on the number of *Striga* attachments and sorghum performance (dry shoot weight) was determined for three replicates. First, the Striga attachments per plant data were analyzed by one-way ANOVA in R as they fulfill the basic ANOVA assumptions. To differentiate treatment means that differ significantly from one another, the LSD comparison in the *agricolae* package in R was used. As the sorghum dry shoot weight data were heteroscedastic, the Kruskal-Wallis’s test was used, and to differentiate the treatment means that differ significantly, Dunn’s test was employed. Benjamini-Hockberg’s method for adjusting the p-values for false discovery rate (FDR) is used for both parametric and non-parametric tests. A cut-off value of alpha = 0.05 was used for both analyses to declare statistical significance.

#### Identifying soil traits that affect Striga infection

The influence of the factorial combination of two independent variables: soil sample (D20 and D21) and soil treatment (natural and gamma-irradiated), on the number of *Striga* attachments per unit root dry weight in *Striga*-susceptible sorghum genotype SQR was assessed using generalized linear modeling in R. Gamma-sterilization of the soils led to an increase in root biomass. The root dry weight was normalized across the soils to reduce its confounding effect on *Striga* attachments. Five biological replicates were considered for each treatment. The results of multiple comparisons of the treatments are provided in **Supplementary Tables ST5** and **ST6**. Furthermore, the pair-wise comparison of *Striga* attachment and relative sorghum dry shoot weight in gamma-irradiated and natural soils was performed by student t-test and results are provided in the main document in **Figure 2**.

#### Soil treatment by genotype interaction effect on Striga infection

Considering the observed impact of the D20 soil microbiome on *Striga* root infection for the susceptible sorghum genotype SQR, we further investigated the contribution of the microbial component of soil D20 to *Striga* suppression for four other sorghum genotypes (SRN-39, Framida, Birhan (PSL5061) and Gubiye (P9401)) that are resistant to *Striga* infection. The data were subjected to GLM by considering host genotype (five levels) and soil treatment (two levels) as factors and results of the multiple comparisons are provided in **Supplementary Tables ST5** and **ST6**. At the same time, the pair-wise comparisons were performed by student t-test. The results are shown in **Figure 2**.

#### Generalized Joint Attribute Modeling (GJAM)

Considering that both the host traits (*Striga* attachment, root, and shoot biomass) and the microbiome represent one mutually dependent response to the environment (soil samples (D20 and D21), soil treatment (natural vs. gamma-irradiated) and *Striga* (presence/absence)), we employed a probabilistic model GJAM (*34*) that covers the joint response of the host and the rhizosphere microbiome community. The GJAM approach accounts for the underlying covariance structure, together with the heterogeneous data types that include composition (e.g., bacterial and fungal ASV composition), counts (*Striga* attachment), and continuous variables (root and shoot dry weight of the host). The analysis was performed using R’s GJAM package (*34*). The dimension of the combined fungal and bacterial OTUs was reduced to 103 features for the modeling. The link for the corresponding code for the data analysis is provided in the data availability section of the manuscript.

## Results

### *Striga* infections of sorghum vary across soil samples

Soil microbiome-mediated suppression of soil-borne plant pathogens can be eliminated by sterilization and transplanted via mixing small soil portions (1-10% v/v) into non-suppressive soils (*22*, *35*, *36*). Based on this model, we developed the plug system for studying sorghum-*Striga*-microbiome interactions. The plug system consists of a soil plug (1% v/v) that serves as the inoculum of indigenous soil microbes to colonize the sorghum roots, which then subsequently grow into a standardized sand mix supplemented (or not) with preconditioned *Striga* seeds (**Figure 1A**). As a proof-of-concept, we introduced the GFP-labeled and rifampicin-resistant rhizobacterial strain *Pseudomonas fluorescens* SS101 in natural or gamma-irradiated soil plugs. Results showed that this strain colonized the sorghum roots that outgrow the soil plug with a longitudinal colonization pattern (**Supplementary Figure SF2c**), typically found for introduced rhizobacterial strains and indigenous rhizobacteria (*37*).

**Figure 1.**
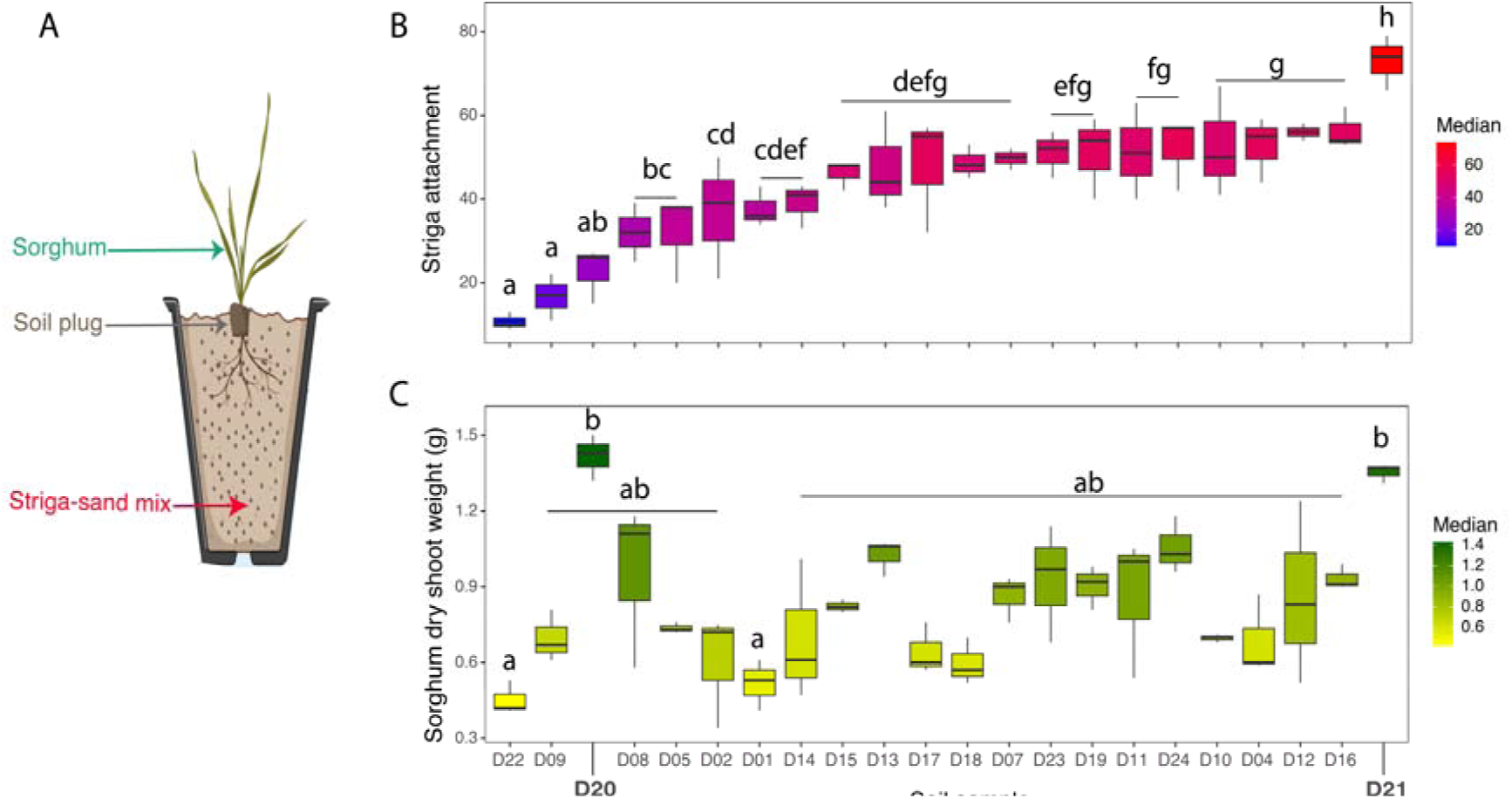
Variation of *Striga* incidence and sorghum performance in 22 soils from (mostly) agricultural field soils in the Netherlands. (**A**) Representation of the plug system (PS) used to screen the variation in *Striga* incidence of sorghum grown in these 22 soils. The soil plug accounts for 1% (v/v) of the potting medium (river sand) and serves as the inoculum of indigenous soil microorganisms; (**B**) *Striga* attachments per plant, and **(C**) sorghum dry shoot weight of the susceptible sorghum genotype ShanQui Red (SQR) grown for six weeks in each of the 22 soils. *Striga* incidence and shoot weight are expressed per plant. Two soil samples (D20 and D21) that showed contrasting *Striga* attachment per plant but with similar sorghum shoot dry weight are marked with larger font sizes. The false color scale shows the median value of Striga attachment and sorghum dry shoot weight. Three biological replicates were considered for each soil. Treatments sharing the same letter are not significantly different.

**Figure 2.**
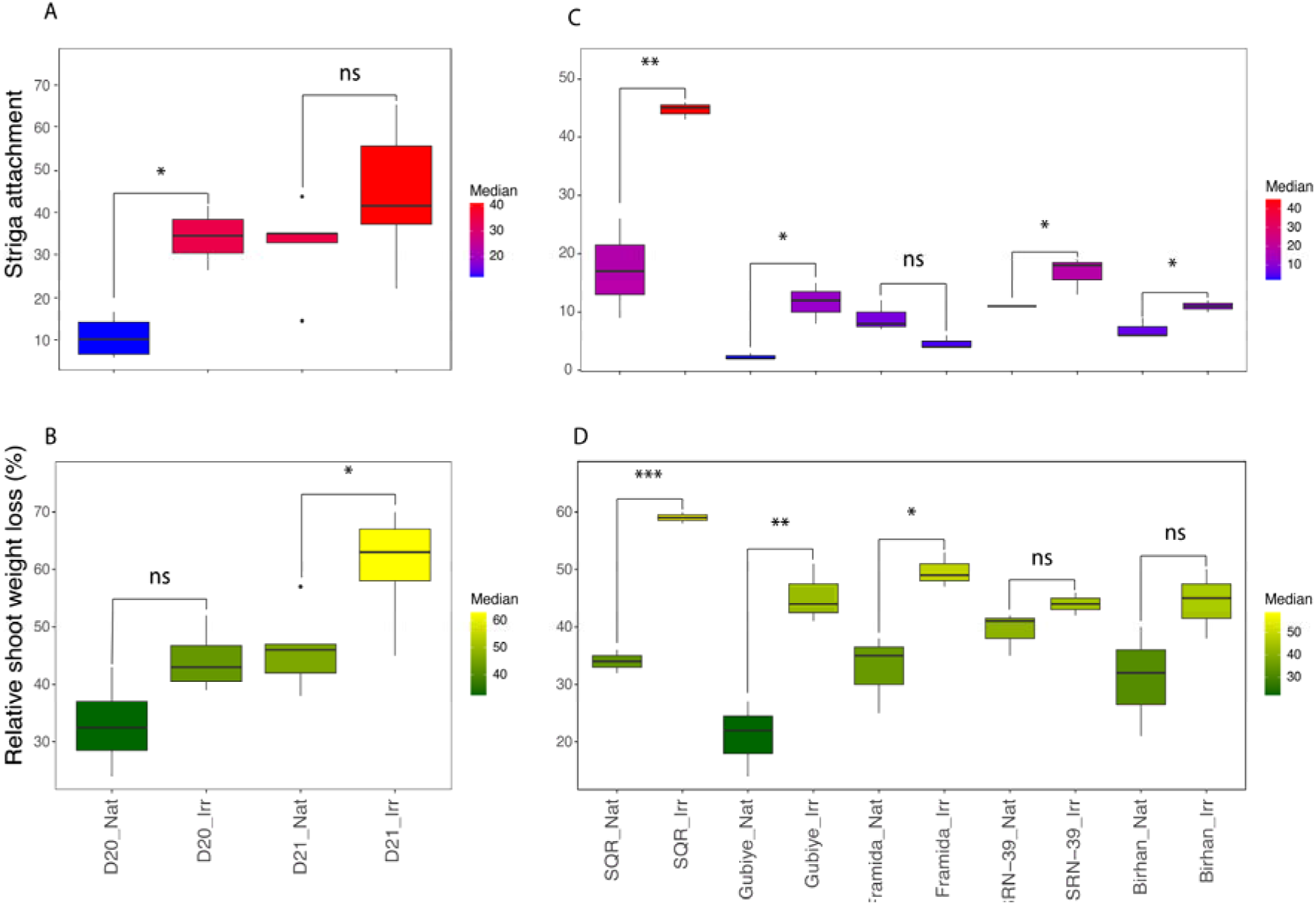
Impact of the soil microbiome on *Striga* incidence and sorghum performance. (**A**) *Striga* attachments per unit root dry biomass, and (**B**) the relative dry shoot weight loss in *Striga*-susceptible sorghum accession ShanQui Red (SQR) compared to the control plants grown in natural (Nat) and gamma-irradiated (Irr) plugs of soils D20 and D21 for six weeks (n=5) in the presence of *Striga*. Shown are (**C**) the number of *Striga* attachments per unit dry root weight, and (**D**) the percent loss in dry shoot weight of four *Striga*-resistant sorghum genotypes (Gubiye (P9401), Framida, and SRN-39, Birhan (PSL5061)) and the *Striga*-susceptible Shan Qui Red (SQR) grown in natural (Nat) and gamma-irradiated (Irr) D20 soil (n=3). The pairwise comparisons were conducted using the student t-test, with statistical significance levels set at *p < 0.05* (\**), p < 0.01 (*\*\*), *p < 0.001 (*\*\*\*), and *p > 0.05* (not significant - ns).

Subsequent screening of the 22 field soils in this plug system showed significant variation (P<0.001) in *Striga* incidence (**Supplementary Table ST2**). The average number of *Striga* attachments per plant ranged from 12 to 73 (**Figure 1B**). Similarly, sorghum showed significant variation (P<0.001) in dry shoot weight (**Supplementary Table ST3**), ranging from 0.5 to 1.3 g (**Figure 1C**). Soils D20 (agricultural) and D21 (pasture soil) showed the lowest and the highest number of *Striga* attachments per plant (*µ* ± SD: 22 ± 6.7 and 73 ±6.6), respectively, without adversely affecting sorghum growth (**Figures 1B** and **C**). Therefore, these two soils, contrasting in *Striga* infection but not in sorghum growth, were used to investigate whether the observed differences in *Striga* infections were associated with the soil microbiome.

### The soil microbiome contributes to *Striga* incidence and sorghum performance in a soil and sorghum genotype-dependent manner

There was no significant interaction effects between the soil samples (D20 or D21) and soil treatments (gamma-irradiated or natural) (P>0.05) on the number of *Striga* attachments per unit of root dry weight (hereafter referred to as *Striga* attachments) (**Supplementary Table ST4**). However, soil sample and treatment showed a significant effect (P<0.001) on the number of *Striga* attachments per unit of root biomass. Consistent with the earlier screening, the *Striga*-susceptible sorghum accession Shan Qui Red (SQR) grown in natural D20 soil showed significantly fewer *Striga* attachments (*µ* ± SD: 12.5 ± 4.8) than SQR grown in soil D21 (*µ* ± SD: 32.4 ± 9.9) (**Figure 2A** and **Supplementary Figure SF3A**). Furthermore, gamma-irradiation significantly increased *Striga* attachments in soil D20 (*µ* ± SD: 33.6 ± 5.5). Also, for soil D21 an increase in *Striga* attachments was observed upon soil sterilization (*µ* ± SD: 43.6 ± 15.5), albeit not statistically significant (P>0.05) (**Figure 2A**). SQR grown in natural D20 soil showed the lowest loss in dry shoot weight (*µ* ± SD: 12.5 ± 4.8) when compared to SQR grown in gamma-irradiated D20 soil (*µ* ± SD: 34.25 ± 6.0) as well as in natural and gamma-irradiated D21 soils (*µ* ± SD: 32.4 ± 9.9 and 43.6 ± 15.5, respectively) (**Figure 2B** and **Supplementary Figure SF3B**).

Considering the apparent impact of the D20 soil microbiome on *Striga* root infection, we further investigated the contribution of the microbiome of soil D20 to *Striga* suppression for four other *Striga*-resistant sorghum accessions (SRN-39, Framida, Birhan (PSL5061) and Gubiye (P9401)); sorghum accession SQR was included as a control. The host genotype x soil treatment interaction effects was significant for *Striga* attachment and the relative dry shoot weight loss (**Supplementary Tables ST5, ST6**). The results show that when grown in natural D20 soil, the number of *Striga* attachments on roots of *Striga*-susceptible SQR (*µ* ± SD: 17.3 ± 8.5) was not significantly different from the number of attachments on roots of the *Striga*-resistant genotypes Framida and SRN-39 (*µ* ± SD: 9.0 ± 2.5 and 11.0 ± 0.0, respectively) (**Supplementary Figure SF3C)**. In gamma-irradiated D20 soil, however, *Striga* incidence increased for SQR (*µ* ± SD: 44.7 ± 1.5) with a significantly higher relative shoot dry weight loss (*µ* ± SD: 59.0 ± 1.0) than the *Striga*-resistant sorghum accessions (**Figures 2C, 2D and Supplementary Figure SF3C and D**). Among the *Striga*-resistant sorghum accessions, gamma-irradiation of the soil led to a significant increase in *Striga* attachments only for Gubiye (*µ* ± SD: from 2.3 ± 0.5 to 11.7± 3.5) and a concomitant loss of dry shoot weight (*µ* ± SD: from 21.0 ± 6.6 to 45.3 ± 5.1). Also, for Framida, a significant loss in dry shoot weight was observed upon gamma-irradiation of the soil (*µ* ± SD: from 32.7 ± 6.8 to 49.7 ± 3.1), but this was not associated with an increase in *Striga* incidence (**Figures 2C** and **2D**). These results suggest that for some but not all *Striga*-resistant sorghum genotypes, the microbiome contributes to *Striga* suppression (cv. Gubiye) or growth (cv. Framida).

### *Striga* infection affects the rhizosphere microbiome composition

Next, we investigated how soil samples, soil sterilization, and the presence of *Striga* influenced the composition of the sorghum rhizosphere microbiome, including fungi (mycobiome) and bacteria (bacteriome), *Striga* attachments, and sorghum growth. To do this, we adopted generalized joint attribute modeling (GJAM) (*34*), a statistical approach that allows us to simultaneously model the relationships between multiple input variables (such as soil sample, sterilization, and *Striga* presence) and various output variables (such as microbiome composition, sorghum growth, and *Striga* attachment). The model estimates the strength and direction of these relationships while accounting for potential interactions between the input and output variables. First, we performed a sensitivity analysis to investigate which input variables or their combinations influenced the output variables. The analysis indicated that a combination of gamma-irradiation of the soil (Irr) and *Striga* presence (+) in soil D21 had the strongest impact on the output variables, followed by irradiation of both soils when accompanied by *Striga* presence. Furthermore, gamma-irradiation of soil D21 and the presence of *Striga* seeds in soil D21 strongly affected the output variables (**Supplementary Figure SF4**). Decomposition of the sensitivity analysis further showed that, compared to the measured host-parasite-related parameters (sorghum root and shoot biomass, shoot weight loss, *Striga* attachments), the effect of treatment combinations was strongest on the rhizosphere microbiome composition (**Figure 3A**).

**Figure 3.**
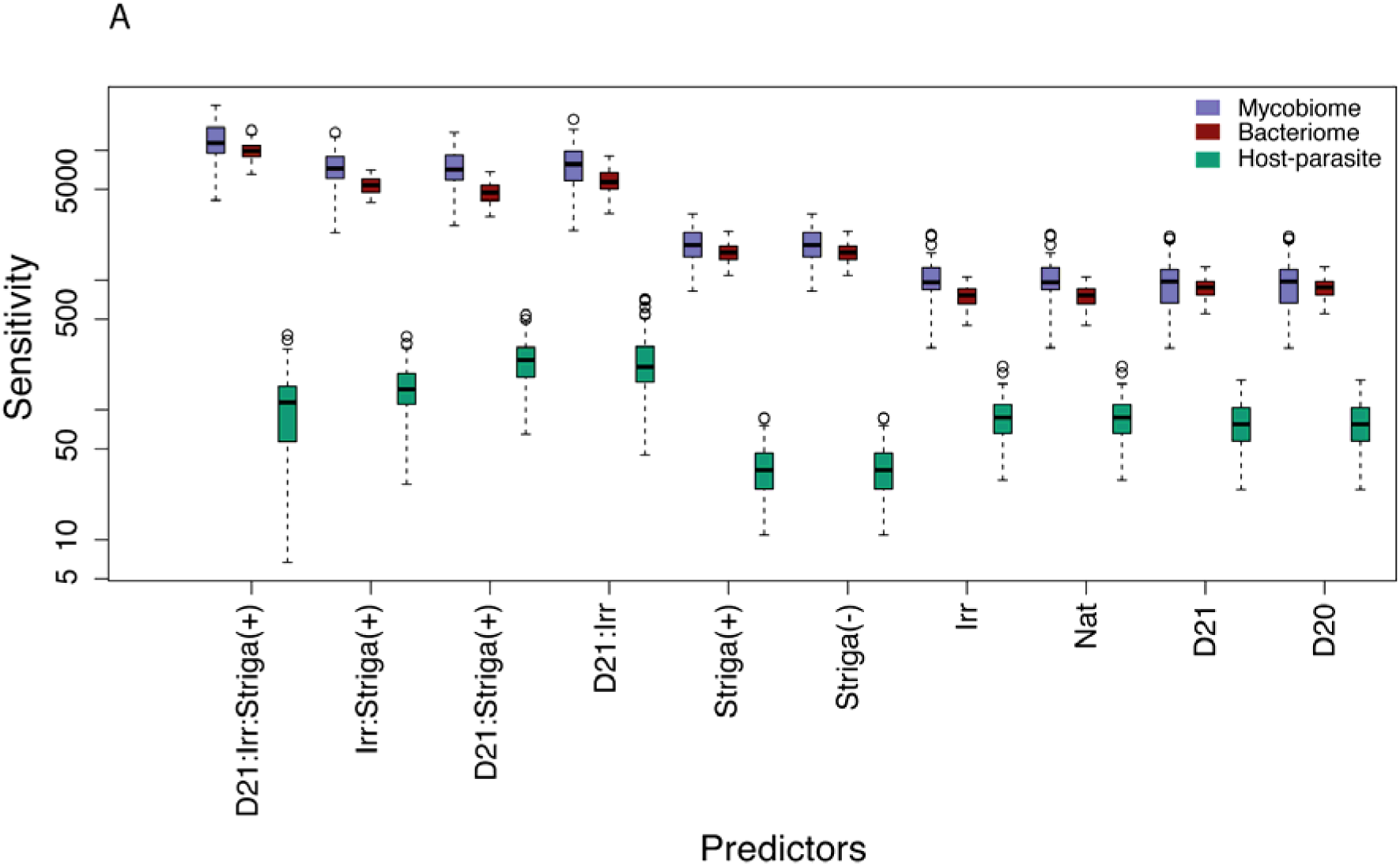
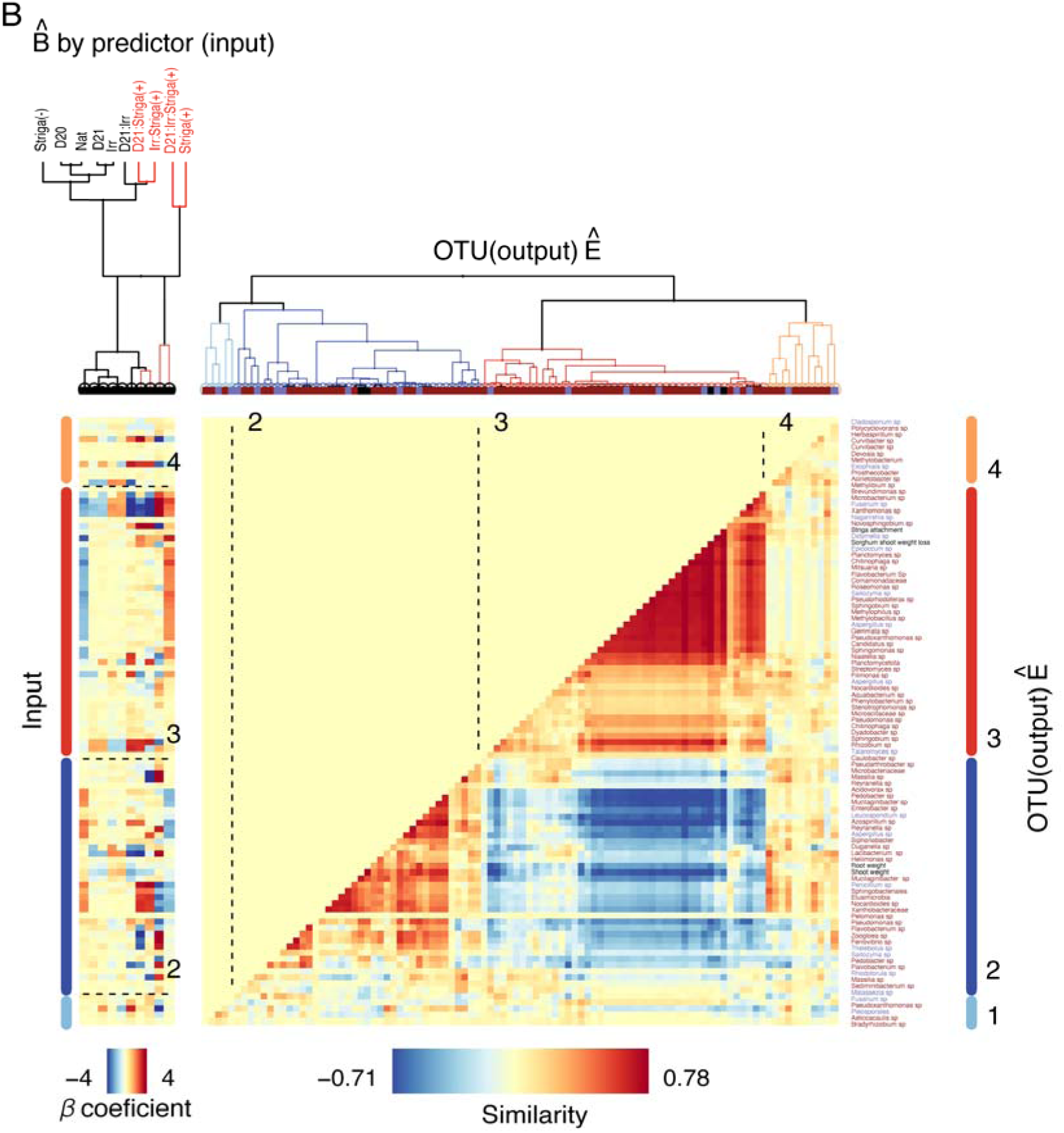
Joint responses of the host and its rhizosphere microbiome to soil samples, soil treatment, and *Striga* presence. (**A**) Sensitivity of host-parasite responses (number of *Striga* attachments, sorghum root and shoot dry weight, sorghum shoot weight loss) and of the rhizosphere microbiome (bacterial and fungal OTUs) to the predictor (factorial combinations of soil samples (D20 and D21), soil treatment (Natural (Nat) and gamma irradiated (Irr)) and *Striga* presence (present (+); absent (-)) in Generalized Joint Attribute Modelling (GJAM). A higher sensitivity indicates the greater importance of the predictor in the model (**B**) Clustering of the most abundant 100 OTUs (97%) and sorghum-associated traits that respond similarly to environmental predictors (input) in GJAM. Response matrix LJ E shows clusters of OTUs based on the similarity of their relationship with environmental predictors using the coefficient matrix LJ B. Warm colors indicate positive, and cool colors indicate negative coefficients. Dotted lines and numbers indicate the four main clusters. Clusters on the left (input variables) and right (output variables) and branches associated with them in the dendrogram (top) were color-coded. Bacterial OUTs are marked brown, fungi blue, and host-associated traits (dry shoot weight, dry root weight, relative dry shoot weight loss, and *Striga* attachment) black in the dendrogram and taxonomy description on the right. The matrix is symmetrical; hence, a mirror image of the matrix is omitted from the top part and colored with a yellow background.

The joint correlation analysis revealed that four clusters and clusters 2 (40 variables) and 3 (45 variables) represent the correlation matrix of most of the total variables in the model (100 variables). The first big cluster (cluster 3) includes bacterial and fungal members of the microbiome with a strong positive correlation with *Striga* attachment, sorghum relative shoot dry weight loss, and environmental factors associated with *Striga* presence and soil sterilization. The mycobiome members were *Alfaria* sp, *Aspergillus* sp, *Didymella* sp, *Epicoccum* sp, *Fusarium* sp, *Nasnishia* sp, and *Talaromyces* sp. Particulary *Epicoccum* sp and *Didymella* sp, *Fusarium* sp, and *Alfaria* sp showed the highest increase in abundance in response to *Striga* presence (**Supplementary Figure SF5**). Intriguingly, these fungal taxa were detected in the sorghum rhizosphere only when plants were challenged with *Striga*. Although several bacterial taxa were present in this cluster, *Mitsuaria chitosanitabida* showed the highest increase in abundance, representing up to ∼40% of the relative frequency of the bacteriome and exhibiting a strong positive correlation with those fungi and *Striga* attachment (**Supplementary Figure SF6**). Also, this cluster represented bacteria classified as *Aeromicrobium*, *Allorhizobium-Neorhizobium-Pararhizobium-Rhizobium*, *Chitinophaga* spp*, Candidatus* sp, Bacterium belonging to the family Comamonadaceae, *Ferrovibrio* sp, *Filimonas* sp, *Flavobacterium* sp, *Hyphomicrobium* sp, *Methylobacillus* sp, *Methylophilus* sp, *Mucilaginibacter* sp, *Niastella* sp, *Ohtaekwangia* sp, *Pantoea* sp, *Pseudorhodoferax* sp, *Pseudoxanthomonas* sp, *Pseudoxanthomonas* sp, *Rivibacter* sp, *Roseomonas* sp, *Sphingobium* spp, and *Sphingomonas* sp were only detected in the rhizosphere when sorghum was challenged with *Striga*. However, compared to the fungi that showed increased abundance when the plant was challenged with *Striga*, the increase in the relative abundance of these bacterial genera was minor (**Figure 3B** **and Supplementary Figures SF7**).

The second large cluster (cluster 2) included members of the microbiome with a strong positive correlation with dry sorghum shoot and root weight and a negative correlation with the presence of *Striga*. *Leucosporidium* sp and *Aspergillus* sp were the two fungi that showed a strong positive correlation with dry sorghum root and shoot weight and the absence of *Striga*. Among the many bacteria, those belonging to the Oxalobacteriacea family are worth mentioning for this Cluster, considering that several of the family members are known for their plant growth-promoting potential. Clusters 2 and 3 showed a strong negative correlation, as shown by the ‘cold’ color in the middle of the correlation matrix (**Figure 3B**).

### The *Striga* seed - sorghum rhizosphere axis

To investigate the impact of the environmental factors on the composition and relatedness of the microbial communities, we performed Beta diversity analysis using Bray-Curtis and Weighted UniFrac distance matrices, respectively. The results revealed that *Striga* had the strongest effect on bacteriome and mycobiome composition among the environmental factors, with significant differences observed in both Bray-Curtis and Weighted UniFrac distances (**Supplementary Table ST7**). Soil samples significantly affected the bacteriome composition as measured by Bray-Curtis, but not the mycobiome composition measured by either metric. Soil treatment did not significantly affect the bacteriome and mycobiome compositions.

Considering the change in the diversity and number of bacterial and fungal species in the sorghum rhizosphere in response to *Striga* presence, we hypothesized that the rhizosphere microbiome of sorghum is partially determined by *Striga* infections or by the microbiome of *Striga* itself. To further investigate if the *Striga* microbiome contributes to sorghum rhizosphere microbiome composition, we performed amplicon sequencing of both the epiphytic and endophytic bacterial and fungal communities of the *Striga* seeds used in the bioassays. Furthermore, we isolated several epiphytes and endophytes from the *Striga* seeds and characterized these taxonomically.

Analyses of the bacterial community composition of the *Striga* seeds revealed that the episphere harbors diverse taxonomic groups, with *Pseudomonas psychrotolerance* dominating the community (27% ± 2 of the reads). This bacterium was the only one detected in the *Striga* seed endosphere (**Figure 4A**). No bacterial OTUs from the *Striga* seeds were detected in the sorghum rhizosphere upon exposure of sorghum roots to *Striga* (**Supplementary Figure SF6**). Some bacterial taxa showed presence in both *Striga* seed and the sorghum rhizosphere, but this was independent of the *Striga* presence or absence.

**Figure 4.**
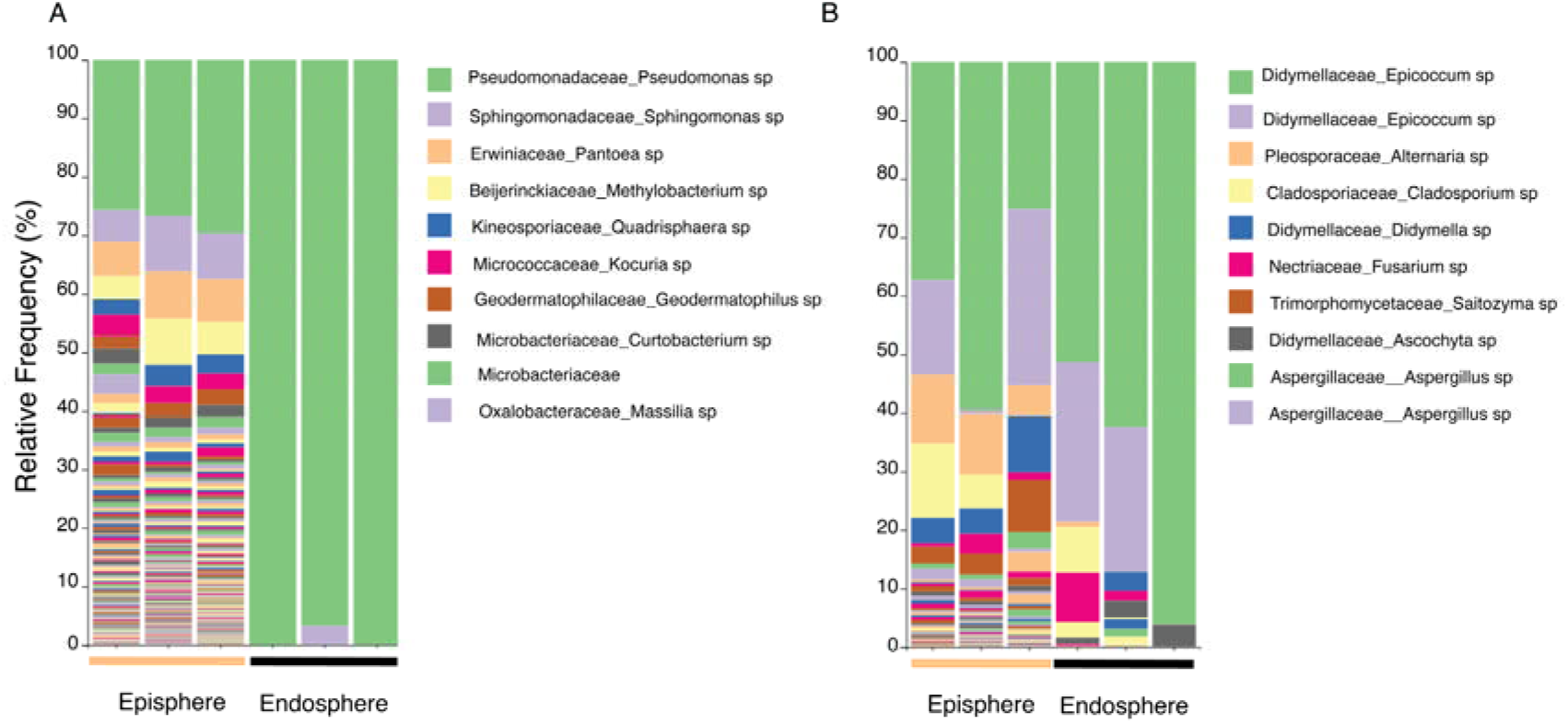
The composition of the epiphytic and endophytic bacteriome (A) and mycobiome (B) of the *Striga* seeds used in the bioassays. The Y-axis depicts the relative abundance of the OTUs for three replicate *Striga* seed batches. The top 10 most abundant bacterial and fungal genera are specified.

The *Striga* seed episphere was also shown to harbor a more diverse fungal community than the endosphere (**Figure 4B**). Nevertheless, *Epicoccum* sp dominated the seed endosphere (70% ± 22 of the reads) and seed episphere (40% ± 17 of the reads). In contrast, *Didymella* sp and *Fusarium* sp were among the fungi detected in the seed endosphere. Culture-based isolation confirmed the abundant presence of *Epicoccum* spp. (*Epicoccum sorghinum* and *Epicoccum pimprinum*), *Didymela* spp. (*Didymella americana* and *Didymella glomerata*), *Fusarium* spp. (*Fusarium sudanense*, *Fusarium nygamai*, *Fusarium andiyazi* and *Fusarium thapsinum*), and *Aspergillus* spp. (*Aspergillus terrus* and *Aspergillus flavus*) in the seed endosphere and episphere. It should be mentioned that *Epicoccum* sp proved to be the dominant taxon in the *Striga*-infested sorghum rhizosphere representing ∼40% of the mycobiome (**Supplementary Figure SF5**). Similarly, most of the other dominant fungal taxa present on/in the *Striga* seeds were detected in the rhizosphere of *Striga*-infested plants (**Supplementary Figure SF5**).

The results further show that the amplicon sequences of the dominant fungal taxa found in both the rhizosphere of *Striga*-infested plants and on/in the *Striga* seeds match the ITS sequences of the fungal isolates obtained from the *Striga* seeds (**Supplementary Table ST8** and **ST9** [https://github.com/DesalegnE/Reciprocal-interactions_Etalo-et-al-2023-]). Subsequent species-level identification of these isolates was conducted using multigene markers. Based on these results, the dominant fungi in the rhizosphere and on/in *Striga* seeds were classified as *Epiccocum sorghinum* (LLC1238), *Fusarium sudanense* (LLC947), *Didymella americana* (LLC3995), and *Alfaria humicola* (LLC985).

## Discussion

With the current developments in microbiome research, it is becoming evident that the plant and the microbiome of different plant tissues form a holobiome, potentially acting as a unit of selection (*38*). Hence, the microbiome can add to plant host genetic variability yet is often neglected in plant breeding strategies. Notably, this is crucial in plant-parasite interactions that occur predominantly belowground, where taxonomically and functionally diverse microbes are abundantly present and potentially contribute to the outcome of those interactions. Here, we investigated the potential of the soil microbiome to interfere with the interaction between the parasitic weed *Striga* and its host, sorghum. We showed significant variation in *Striga* attachments to roots of sorghum grown in 22 different field soils. Furthermore, gamma-irradiation of the most *Striga*-suppressive soil led to a substantial increase in *Striga* incidence not only for the susceptible sorghum genotype SQR but, at least to some extent, also for one of the *Striga*-resistant sorghum genotypes.

The 22 tested soils varied considerably in microbiome composition and physicochemical properties (**Supplementary Figure SF1**). The *Striga*-susceptible sorghum accession SQR grown in these soil plugs showed a 6-fold variation in the number of *Striga* attachments and up to 3-fold differences in sorghum shoot dry weight. It should be noted that many soils that exhibited relatively low *Striga* attachments also showed reduced sorghum performance (**Figure 1**). This was not the case for soil D20, with low numbers of *Striga* attachments and good sorghum performance. The *Striga-*suppressive effect conferred by the D20 soil microbiome to the susceptible sorghum genotype SQR was striking and comparable to that found for the *Striga*-resistant sorghum genotypes SRN-39, Framida, and Birhan. Yet, some sorghum genotypes (Birhan, SRN-39) suppressed *Striga* attachment independently of the soil microbiome, while genotype Gobiye partially relies on the microbiome to suppress *Striga* infection. It is tempting to speculate that for this latter sorghum genotype, breeding for *Striga* resistance may have unintentionally co-selected for plant trait(s) that recruit(s) microbiome members that counteract *Striga* infection. Irregular durability and stability of sorghum resistance to *Striga* across different locations is the major setback in the resistance breeding (*39–41*). The prevalence of genetically diverse *Striga* ecotypes was suggested to be a major reason for this inconsistent *Striga* suppressive effect of resistant genotypes in different agroecological zones (*40*, *41*). Our results suggest potential microbiome-mediated contributions to the observed inconsistencies of resistant sorghum accessions in suppressing *Striga* infection in different field soil locations.

Our result further showed that the soil microbiome is critical for suppressing *Striga* attachments. Although the number of *Striga* attachments is significantly lower in resistant sorghum accessions, these few infections resulted in significant weight loss (∼20-50%) of the sorghum shoots in a genotype-dependent manner. This is evident for Gobiye and Framida, that showed a significant increase in shoot weight loss when the soil is gamma irradiated, suggesting that the microbiome contributes to *Striga* tolerance. Identifying the genetic bases of resistance and tolerance in light of the microbiome could be an important step for the future breeding of resilient crops against parasitic weeds.

Considering the significant effects of the soil microbiome on *Striga* infection and relative yield loss, we profiled both bacterial and fungal rhizosphere communities of the susceptible sorghum genotype grown in natural and gamma-irradiated soils D20 and D21 in the presence and absence of *Striga*. Using probabilistic modeling (GJAM) (*34*), we analyzed the impact of individual predictors and their combinations of different variables. GJAM is a robust tool to integrate multivariate, multiple scales and a mixture of continuous and discrete observations. Such versatility enabled us to track the microbial community (compositional data) and host and parasite traits (shoot & root dry weight, relative shoot weight loss, and *Striga* attachment). The sensitivity analysis revealed that the presence of *Striga* had the highest impact on the variables in the conducive natural soil (D21) and both the conducive (D21) and suppressive (D20) soils when they were gamma-irradiated. These three conditions also showed a marked increase in *Striga* attachment and sorghum shoot weight loss compared to plants grown in natural D20 soil with an intact microbiome (**Figures 3A** and **B**). The impact on the rhizosphere microbiome was the highest compared to variables corresponding to the host and the parasite performance. This suggests that modulation of the rhizosphere microbiome is a key determinant of *Striga* infection and the concomitant impact on host performance.

To dissect the relationship between host performance, *Striga* attachment, and rhizosphere microbiome, we performed a joint correlation analysis of these variables considering the influencing soil environmental factors. The analysis revealed a clear correlation structure among the fungal and bacterial members of the microbiome with *Striga* attachment and sorghum relative shoot weight loss. Particularly the high positive correlation between *Mitsuaria chitosanitabida* and other bacteria belonging to *Chitinophaga* genera, *Flavobacterium* sp, with the predominant fungi that colonize the host rhizosphere, such as species of *Didymella*, *Epicoccum*, and *Fusarium* is intriguing. These groups of bacteria can solubilize chitin, an integral structural component of fungi, and are often observed to be enriched in response to fungal invasion of the rhizosphere (*42–44*). Moreover, the Beta diversity analysis also showed that *Striga’s* presence significantly changed the sorghum rhizosphere microbiome composition. This might suggest a potential change in rhizosphere microbial function in response to *Striga* infection, likely determining the interaction’s outcome, hallmarking the interaction’s tripartite nature. Interactions between *Striga* and the rhizosphere microbiota are likely as they share the same environment and compete for host-derived resources. Supporting this view, parasites from different phyla showed host-microbiome composition modulatory effects (*45–47*). Still, it is unknown if the modulation is beneficial for the parasite, represents a defense mechanism of the host, or is the consequence of parasite-mediated alteration of host metabolism (e.g., root exudation) that affects the composition of the rhizosphere microbiome. Furthermore, if the change in microbiome structure leads to a change in the associated microbial functions favoring the parasite remains to be investigated.

Microbiome profiling of the *Striga* seed episphere and endosphere showed that several *Striga* seed-associated fungi (SAF) are found in the sorghum rhizosphere only in the presence of *Striga*. How they colonize the sorghum rhizosphere and if they have a role in the *Striga* infection process remains to be investigated. Interestingly, some of these SAF have been reported as sorghum pathogens, suggesting a potential co-evolutionary relationship between sorghum pathogens and *Striga* (*48–51*). Several of these SAF are also shown to produce secondary metabolites with antibacterial activities (*52–54*). Hence, the SAF may also determine *Striga’s* observed and marked impact on the bacterial community composition of the sorghum rhizosphere. This remains speculative and requires an elaborate series of new experiments to pinpoint the importance and consequences of these interactions between the microbiomes of the parasite and the sorghum rhizosphere.

It should be noted that the amplicon sequencing used in our research provided limited information on the spatiotemporal dynamics of the root microbiome and their associated functions that are relevant to the suppression of *Striga*. Therefore, future studies should include metagenomics, metatranscriptomics, and metabolomics to shed light on enriched biosynthetic gene clusters and their corresponding microbial enzymes and metabolites of the active members of the microbiome community in *Striga*-suppressive soils. This is crucial in designing a versatile synthetic microbial community and soil amendment strategies to enrich targeted microbial functions to suppress *Striga* and improve sorghum growth.

## Authors’ contributions

JMR and DWE conceived the project; AO performed collection and physicochemical analysis of the soils; TT performed collection and characterization of the *Striga* seeds and Sorghum genotypes; DWE & DR performed the experiments; LL and PWC performed the isolation and identification of the Striga seed associated fungal strains; DWE, FDA, MFAL, and EK analyzed the data; DWE and JMR wrote the manuscript with input and contributions of all authors.

## DATA AVAILABILITY

The raw reads supporting this study’s findings have been deposited in the NCBI BioProject database (https://www.ncbi.nlm.nih.gov/bioproject/) with links to BioProject accession numbers PRJNA982817. The codes used to process the data can be found at (https://github.com/DesalegnE/Reciprocal-interactions_Etalo-et-al-2023-).

