## Supplementary Table ST1-ST8 for "Reciprocal interactions between the sorghum root microbiome and the parasitic weed *Striga hermonthica*"

| **Supplementary Table ST1.** Physicochemical characteristics and cropping story of soils collected from predominantly agricultural fields in the Netherlands. | | | | | | | | | | | | | | | | |
| --- | --- | --- | --- | --- | --- | --- | --- | --- | --- | --- | --- | --- | --- | --- | --- | --- |
| Soil | pH | AVR Fe | AVR K | AVR Mg | AVR P | AVR S | AVR OM | AVR C [%] | AVR N [%] | AVR C/N [%] | type | Organic | crop 2016 | crop 2015 | crop 2014 | Coordinates |
| D1 | 6.22 | 0.09 | 34.00 | 163.19 | 1.40 | 1.29 | 3.46 | 2.31 | 0.14 | 16.55 | Sand | N | N | N | N | 51.543654761305135 , 5.8517040686788535 |
| D2 | 6.75 | 0.03 | 86.66 | 36.86 | 0.89 | 2.50 | 3.81 | 1.60 | 0.11 | 15.04 | Clay | N | Wheat | Potato | Wheat | 51.76116461190135 , 4.4217229869102646 |
| D4 | 7.3 | 0.02 | 117.12 | 56.49 | 1.53 | 7.21 | 3.10 | 2.11 | 0.13 | 16.77 | Clay | Y | Wheat | Onion | Lucerne | 52.67733000460026 , 5.845727409204449 |
| D5 | 6.47 | 0.16 | 109.72 | 74.87 | 7.76 | 1.29 | 3.56 | 1.99 | 0.16 | 12.73 | Clay | N | Wheat | Potato | Wheat | 51.34332701656123 , 3.6061494684038937 |
| D7 | 7.58 | 0.03 | 82.92 | 61.11 | 1.66 | 5.57 | 4.32 | 2.92 | 0.18 | 16.08 | Sand | N | Wheat | Potato | Wheat | 53.33042679058607,6.275790569493179 |
| D8 | 6.87 | 0.08 | 181.54 | 109.68 | 16.23 | 1.14 | 2.81 | 1.94 | 0.16 | 12.47 | Sand | Y | Wheat | Yakon | Yakon | 51.27258828910626,6.0062516036353975 |
| D9 | 6.87 | 0.06 | 174.31 | 113.01 | 3.06 | 1.26 | 3.26 | 1.35 | 0.12 | 11.40 | Sand | Y | Wheat | Onion | Corn | 51.272120763051774,6.008179006262658 |
| D10 | 7.49 | 0.02 | 97.07 | 70.88 | 1.12 | 2.96 | 2.92 | 2.62 | 0.16 | 16.65 | Clay | Y | Spelt | Pea | Beetroot, pumpkin | 51.34672350770874,3.582965038353318 |
| D11 | 7.28 | 0.11 | 68.77 | 56.43 | 5.43 | 1.17 | 3.48 | 1.49 | 0.16 | 12.65 | Sand | Y | Luzerne | Luzerne | Barley | 51.54839951928703,4.3042692532241205 |
| D12 | 7.75 | 0.02 | 233.38 | 83.72 | 1.06 | 9.00 | 3.81 | 2.03 | 0.14 | 14.37 | Clay | Y | Wheat | Beetroot | Potato | 51.77042075724939,4.493490193385289 |
| D13 | 7.55 | 0.04 | 207.13 | 122.52 | 3.40 | 7.57 | 7.03 | 3.82 | 0.34 | 11.30 | Sand/Clay | Y | Grass | Grass | Cabbage | 51.770778278187606,4.487125277375227 |
| D14 | 7.61 | 0.04 | 82.49 | 98.91 | 0.70 | 8.61 | 5.04 | 2.35 | 0.15 | 15.31 | Clay | N | Wheat | Beetroot | Potato | 52.45982040396538,5.637314757295926 |
| D15 | 7.82 | 0.02 | 45.83 | 60.60 | 0.90 | 5.95 | 3.77 | 1.91 | 0.11 | 16.84 | Clay | Y | Potato | Grass | Onion | 52.54409238258504,5.5807603597519755 |
| D16 | 7.63 | 0.03 | 53.60 | 78.47 | 5.61 | 2.79 | 2.91 | 1.50 | 0.10 | 14.53 | Clay | Y | Luzerne | Potato | Carrot | 51.65545377639545,4.4860969515006355 |
| D17 | 7.75 | 0.01 | 87.17 | 74.54 | 1.03 | 2.27 | 4.23 | 2.63 | 0.15 | 17.82 | Clay | Y | Potato | Spelt | Pumpkin | 51.346005752252395,3.581139420903252 |
| D18 | 7.69 | 0.02 | 53.59 | 113.54 | 1.49 | 5.80 | 7.53 | 2.30 | 0.24 | 9.65 | Clay | N | Wheat | Wheat | Wheat | 53.18724569478442,7.175740124145935 |
| D19 | 7.73 | 0.02 | 63.09 | 104.28 | 1.50 | 5.00 | 5.38 | 2.35 | 0.24 | 9.70 | Clay | N | Wheat | Wheat | Sugar_beet | 53.18602861507295,7.175716884873125 |
| D20 | 7.3 | 0.10 | 234.45 | 299.63 | 2.69 | 6.98 | 8.00 | 2.88 | 0.36 | 9.37 | Sand | N | Maize | Maize | Wheat | 51.97487050892559,6.205089101755299 |
| D21 | 5.56 | 0.82 | 58.82 | 484.44 | 4.86 | 24.70 | 20.59 | 9.30 | 1.04 | 8.92 | Sand | N/A | Pasture | Pasture | Pasture | 51.92733006262468,5.092160040511496 |
| D22 | 6.5 | 0.03 | 74.84 | 237.38 | 0.87 | 3.67 | 3.02 | 2.65 | 0.30 | 8.71 | Clay | N | Maize | Wheat | Sugar_beet | 51.75145259676526,5.15843045368626 |
| D23 | 5.92 | 0.47 | 51.98 | 122.23 | 11.88 | 4.03 | 4.50 | 1.92 | 0.17 | 11.07 | Sand | N | Maize | Maize | Maize | 51.71172009291613,5.492069152170564 |
| D24 | 7.27 | 0.02 | 76.47 | 66.70 | 2.58 | 5.24 | 6.15 | 0.98 | 0.11 | 8.88 | Clay | N | Wheat | Beetroot | Wheat | 50.972230157625525,5.959331933190798 |

**Supplementary Table ST2** (Soil sample on *Striga* attachment).

| Anova Table (Type III tests) | | | | |
| --- | --- | --- | --- | --- |
| Response: Striga attachment | | | | |
|  | Sum Sq | Df | F value | Pr(>F) |
| (Intercept) | 4256.3 | 1 | 61.9308 | 6.213e-10 *** |
| Soil sample | 12929.0 | 21 | 8.9581 | 7.036e-10 *** |
| Residuals | 3024.0 | 44 |  |  |
| Signif. codes: 0 ‘***’ 0.001 ‘**’ 0.01 ‘*’ 0.05 ‘.’ 0.1 ‘ ’ 1 | | | | |

| **Supplementary Table ST3** (Soil sample on Sorghum performance). | | | | |
| --- | --- | --- | --- | --- |
| Anova Table (Type III tests) | | | | |
| Response: Sorghum shoot dry weight | | | | |
|  | Sum Sq | Df | F value | Pr(>F) |
| (Intercept) | 1.0302 | 1 | 33.3442 | 3.199e-07 *** |
| Soil | 3.9411 | 21 | 6.0741 | 2.427e-08 *** |
| Residuals | 1.7920 | 58 |  |  |
| Signif. codes: 0 ‘***’ 0.001 ‘**’ 0.01 ‘*’ 0.05 ‘.’ 0.1 ‘ ’ 1 | | | | |

**Supplementary Table ST4** (Soil sample x soil treatment on *Striga* attachment).

| Anova Table (Type II tests) | | | | |
| --- | --- | --- | --- | --- |
| Response: Striga attachment per unit root dry root weight | | | | |
|  | Sum Sq | Df | F value | Pr(>F) |
| Soil sample (SS) | 770.06 | 1 | 27.2790 | 0.0002137 *** |
| Soil treatment (STR) | 663.06 | 1 | 23.4886 | 0.0004006 *** |
| SS:STR | 68.06 | 1 | 2.4111 | 0.1464438 |
| Residuals | 338.75 | 12 |  |  |
| Signif. codes: 0 ‘***’ 0.001 ‘**’ 0.01 ‘*’ 0.05 ‘.’ 0.1 ‘ ’ 1 | | | | |

**Supplementary Table ST5** (Genotype x soil treatment on Sorghum performance).

| Analysis of Deviance Table (Type II tests) | | | |
| --- | --- | --- | --- |
| Response: Striga attachment per unit root dry root weight | | | |
|  | LR Chisq | Df | Pr(>Chisq) |
| Genotype (G) | 155.278 | 4 | 2.2e-16 *** |
| Soil treatment (STR) | 43.708 | 1 | 3.813e-11 *** |
| G:STR | 26.405 | 4 | 2.622e-05 *** |
| Signif. codes: 0 ‘***’ 0.001 ‘**’ 0.01 ‘*’ 0.05 ‘.’ 0.1 ‘ ’ 1 | | | |

**Supplementary Table ST6** (Genotype x soil treatment on Sorghum performance).

| Analysis of Deviance Table (Type II tests) | | | |
| --- | --- | --- | --- |
| Response: Percent loss in shoot dry weight | | | |
|  | LR Chisq | Df | Pr(>Chisq) |
| Genotype (G) | 20.152 | 4 | 0.0004661 *** |
| Soil treatment (STR) | 48.413 | 1 | 3.453e-12 *** |
| G:STR | 15.889 | 4 | 0.0031719 ** |
| Signif. codes: 0 ‘***’ 0.001 ‘**’ 0.01 ‘*’ 0.05 ‘.’ 0.1 ‘ ’ 1 | | | |

**Supplementary Table ST7.** Beta diversity analysis based on Bray Curtis (composition) and Weighted UniFrac (phylogeny).

| Environmental factor | Sample size | Beta diversity | | | |
| --- | --- | --- | --- | --- | --- |
|  |  | Bray-Curtis (composition) | | Weighted UniFrac (phylogeny) | |
|  |  | Bacteriome | Mycobiome | Bacteriome | Mycobiome |
|  |  | q-value | q-value | q-value | q-value |
| Striga | 39 | 0.001* | 0.001* | 0.001* | 0.001* |
| Soil sample | 39 | 0.01* | 0.306 | 0.003* | 0.238 |
| Soil treatment | 39 | 0.156 | 0.369 | 0.074 | 0.369 |

*Striga* (+, -), Soil sample (D20, D21), Soil treatment (Natural, irradiated), Permutation = 999

**Supplementary Table ST8.** Sequence similarity between ITS partial sequences of dominant fungi from sorghum rhizosphere and *Striga* seeds with Full length ITS sequence of Fungi isolated from *Striga* seed.

| ***Epicoccum sorghinum*** | | | | | | | |
| --- | --- | --- | --- | --- | --- | --- | --- |
| Sample | OUT (Query) | Subject (Full ITS) | Subject source | Score | Identities (Query length) | Percentage | Expect |
| Seed (partial ITS1) | OUT_01 | LLC1238 | *Striga* seed isolates ITS sequence | 416 | 250/250 (284) | 100 | 3.00E-129 |
|  | OUT_02 | LLC1238 | *Striga* seed isolates ITS sequence |  | 251/251 (284) | 100 | 9.00E-130 |
| Rhizosphere (partial ITS2) | OUT_02 | LLC1238 | *Striga* seed isolates ITS sequence | 369 | 204/204 (204) | 100 | 1.00E-104 |
| ***Fusarium sudanense*** | | | | | | | |
| Seed (partial ITS1) | OUT_15 | LLC947 | *Striga* seed isolates ITS sequence | 453 | 251/251 (277) | 100 | 5.00E-130 |
| Rhizosphere (partial ITS2) | OUT_03 | LLC947 | *Striga* seed isolates ITS sequence | 340 | 188/188 (188) | 100 | 5.00E-96 |
| ***Didymella americana*** | | | | | | | |
| Seed (partial ITS1) | OUT_06 | LLC3695 | *Striga* seed isolates ITS sequence | 425 | 235/235 (286) | 100 | 2.00E-121 |
| Rhizosphere (partial ITS2) | OUT_05 | LLC3695 | *Striga* seed isolates ITS sequence | 354 | 196/196 (196) | 100 | 2.00E-100 |
| The blast search was performed using Viroblast web server (Deng W, Nickle DC, Learn GH, Maust B, and Mullins JI. 2007. ViroBLAST: A stand-alone BLAST web server for flexible queries of multiple databases and user's datasets. [Bioinformatics 23(17):2334-2336.](http://bioinformatics.oxfordjournals.org/cgi/content/full/23/17/2334)). | | | | | | | |
