## Supplementary Figure SF1-SF7 for "Reciprocal interactions between the sorghum root microbiome and the parasitic weed *Striga hermonthica*"

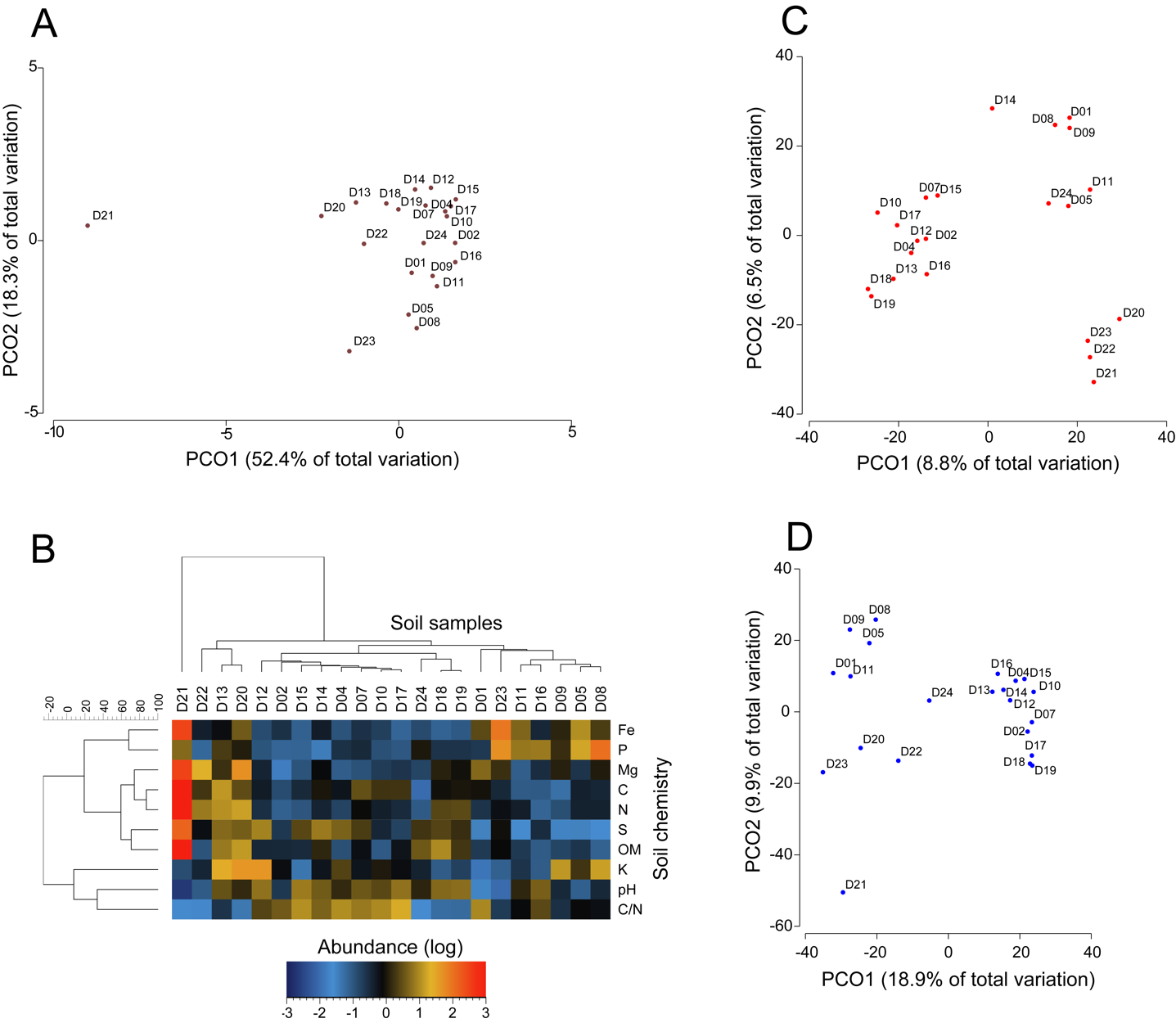

**Supplementary Figure SF1.** Chemical and microbiological profiles of predominantly agricultural soils collected from different parts of the Netherlands. **A**) Principal coordinate analysis (PCoA) depicting variation in soil biochemical profiles between the soil samples. The percent variation explained by the principal coordinates (PCo) are indicated along the x- and y-axis. **B**) Heat map showing the variation in different soil biochemical attributes between the soil samples. **C**) Principal coordinate analysis (PCoA) plot of the bacteria community and **D**) fungi community structures in different bulk soil samples.

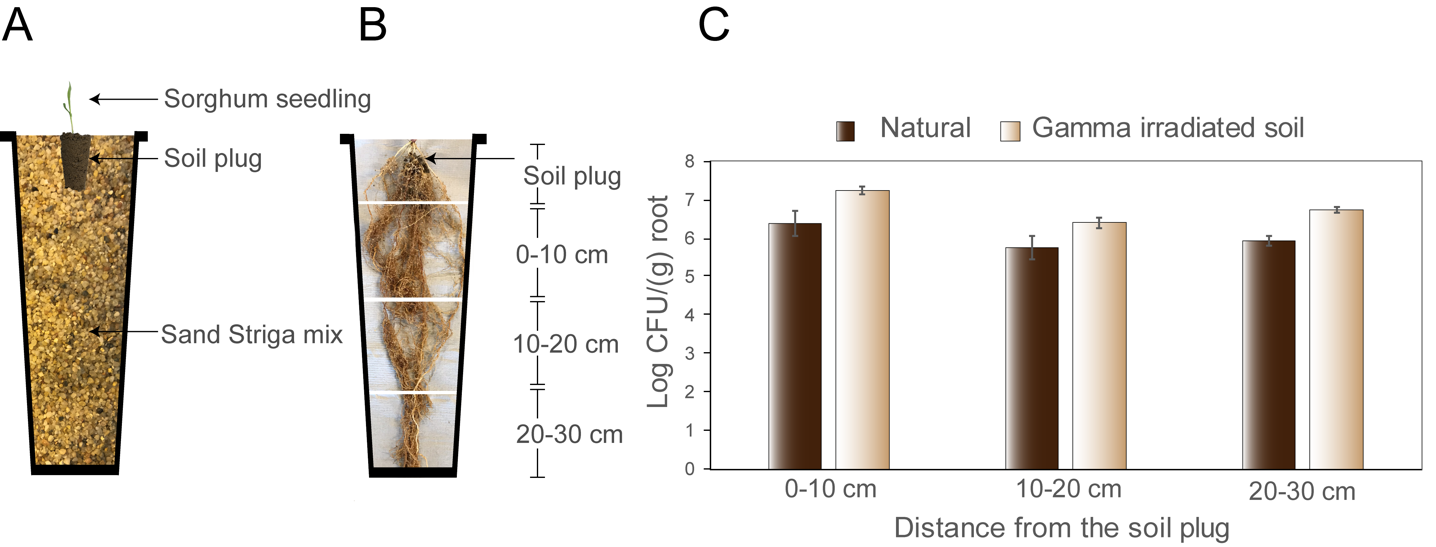

**Supplementary Figure SF2**. The plug system (PS): a tool to study the role of the soil microbiome in parasitic weed-host interaction. (**A**) The PS consists of 5000 germinable *Striga* seeds incorporated into 3 kg fine- and medium-sized sand mixture (1:1) per pot. A 30 g soil plug was embedded at the center of the sand-*Striga* mix, and a pre-germinated sorghum seedling was planted at the plug's center. (**B**) Root sections used to assess the migration of introduced, rifampicin-resistant *Pseudomonas fluorescens* strain SS101 (*Pf* SS101) from the plug to the distal parts of the roots. (**C**) *Pf* SS101 densities associated with the different sections of sorghum roots grown in natural and gamma-irradiated soil plugs (n=3). The experiment was performed in the absence of *Striga*.

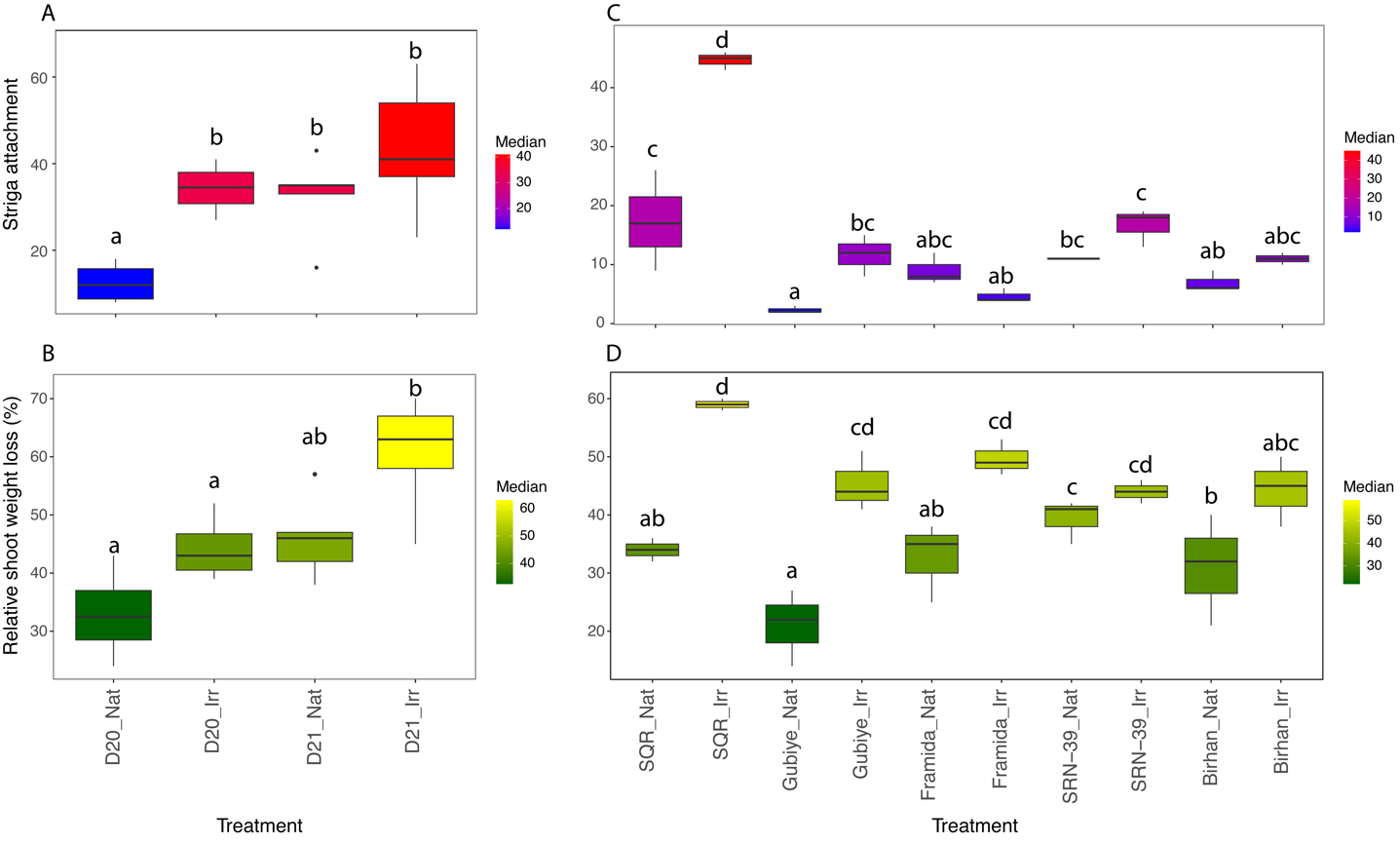

**Supplementary Figure SF3**. Impact of the soil microbiome on *Striga* incidence and sorghum performance. (**A**) *Striga* attachments per unit root dry biomass, and (**B**) the relative dry shoot weight loss in *Striga*-susceptible sorghum accession ShanQui Red (SQR) compared to the control plants grown in natural (Nat) and gamma-irradiated (Irr) plugs of soils D20 and D21 for six weeks (n=5) in the presence of *Striga*. Shown are (**C**) the number of *Striga* attachments per unit root dry weight, and (**D**) the percent loss in dry shoot weight of four *Striga*-resistant sorghum genotypes (Gubiye (P9401), Framida, and SRN-39, Birhan (PSL5061)) and the *Striga*-susceptible Shan Qui Red (SQR) grown in natural (Nat) and gamma-irradiated (Irr) D20 soil (n=3). Treatments sharing the same letters are not significantly different. This figure is a replica of **Figure 2** in the main document, with the only difference being the significance of multiple comparisons displayed across the treatments in SF4.

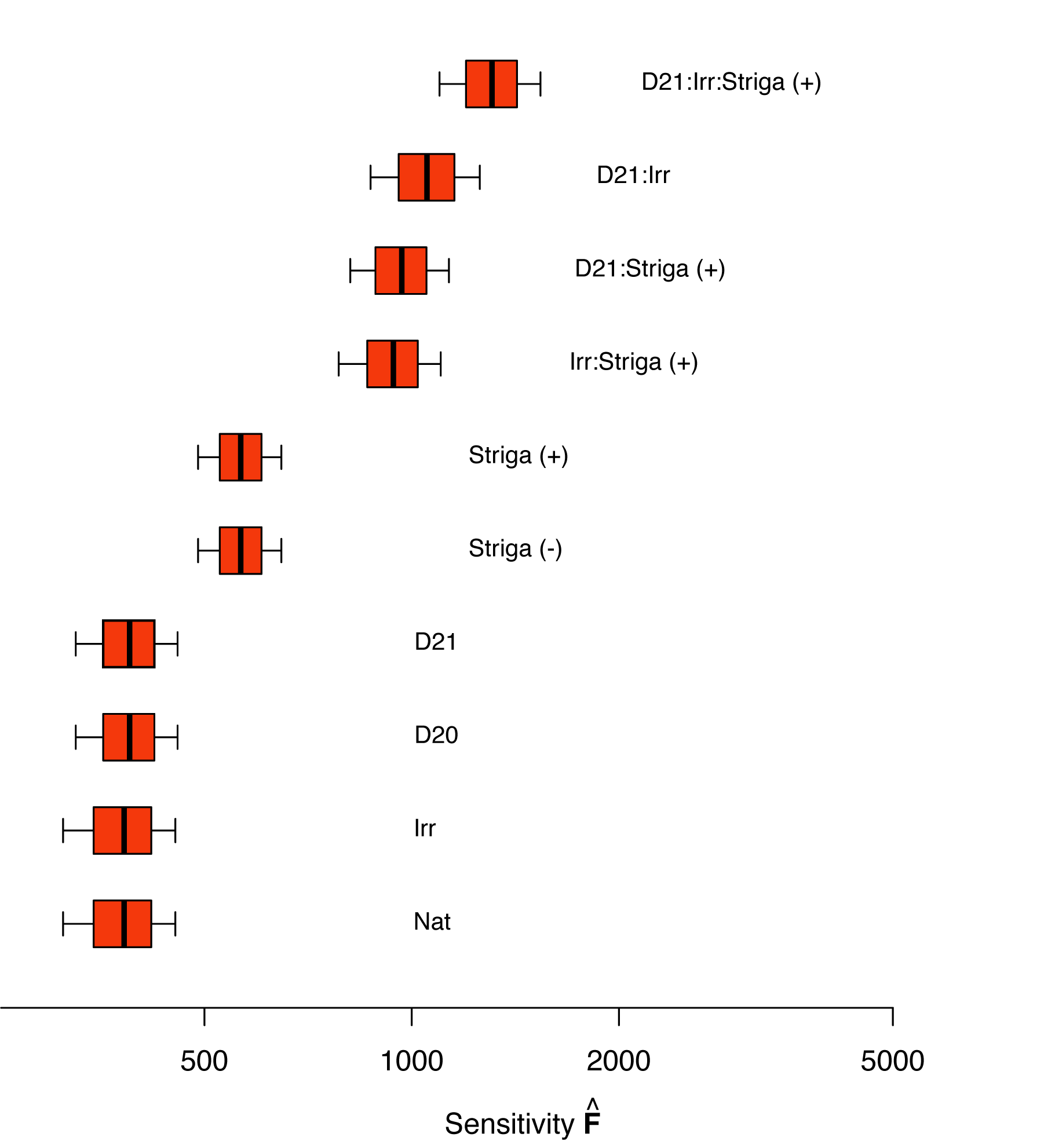

**Supplementary Figure SF4**. Joint responses of sorghum and the rhizosphere microbiome to soil type, soil treatment, and *Striga* pressure. Sensitivity of host-parasite responses (number of *Striga* attachment, sorghum root and shoot dry weight) and the rhizosphere microbiome (bacterial and fungal ASVs) to the environment (soil type (D20/D21), soil treatment (Nat/Irr) and *Striga* pressure ((present (+) and absence (-)). Higher sensitivity indicates the predictors' greater impact on the output variables (the plant and its microbiome).

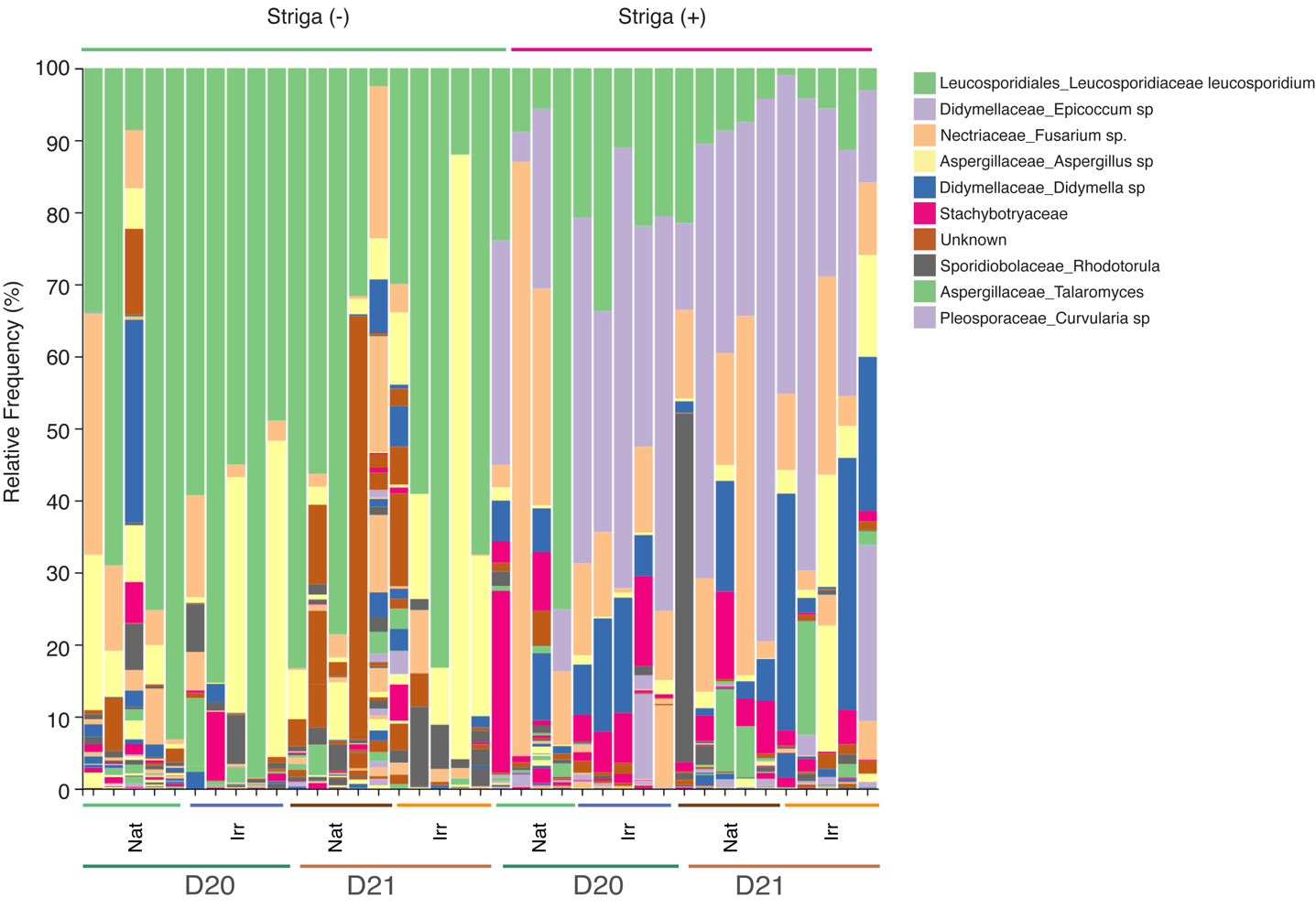

**Supplementary Figure SF5**. Impact of Striga on sorghum rhizosphere mycobiome. Striga susceptible sorghum accession Shan Qui Red grown in natural (Nat) and gamma-irradiated (Irr) portions of two soils (D20 and D21) for six weeks (n=5) in the presence (+) and absence (-) of Striga. The relative abundance of the different taxa is shown in the Y-axis.

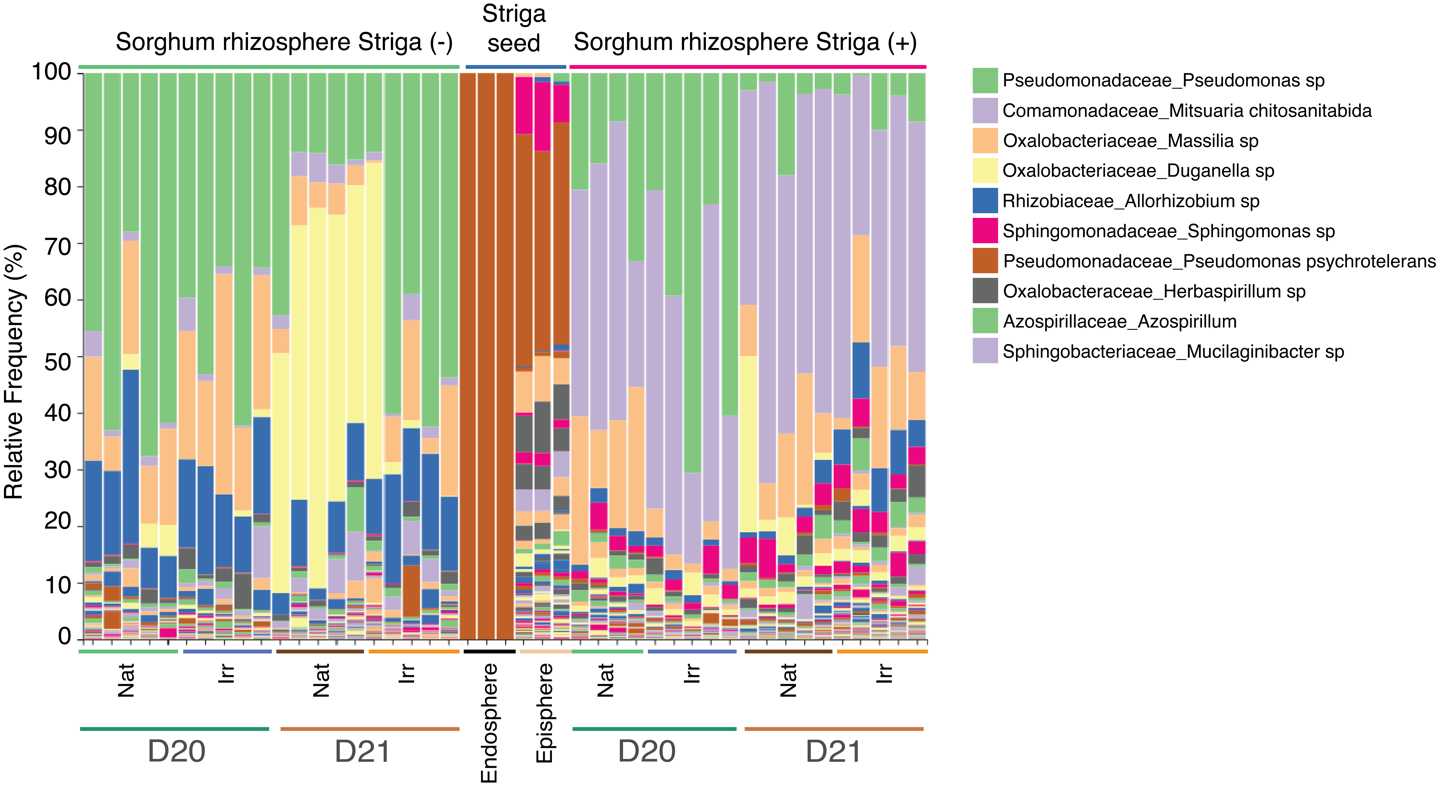

**Supplementary Figure SF6**. Impact of Striga on sorghum rhizosphere bacteriome. Striga susceptible sorghum accession Shan Qui Red grown in natural (Nat) and gamma-irradiated (Irr) portions of two soils (D20 and D21) for six weeks (n=5) in the presence (+) and absence (-) of Striga. The bar plot also shows the episphere and endosphere bacterial community of Striga seed used in the bioassay. The relative abundance of the different taxa is shown in the Y-axis.

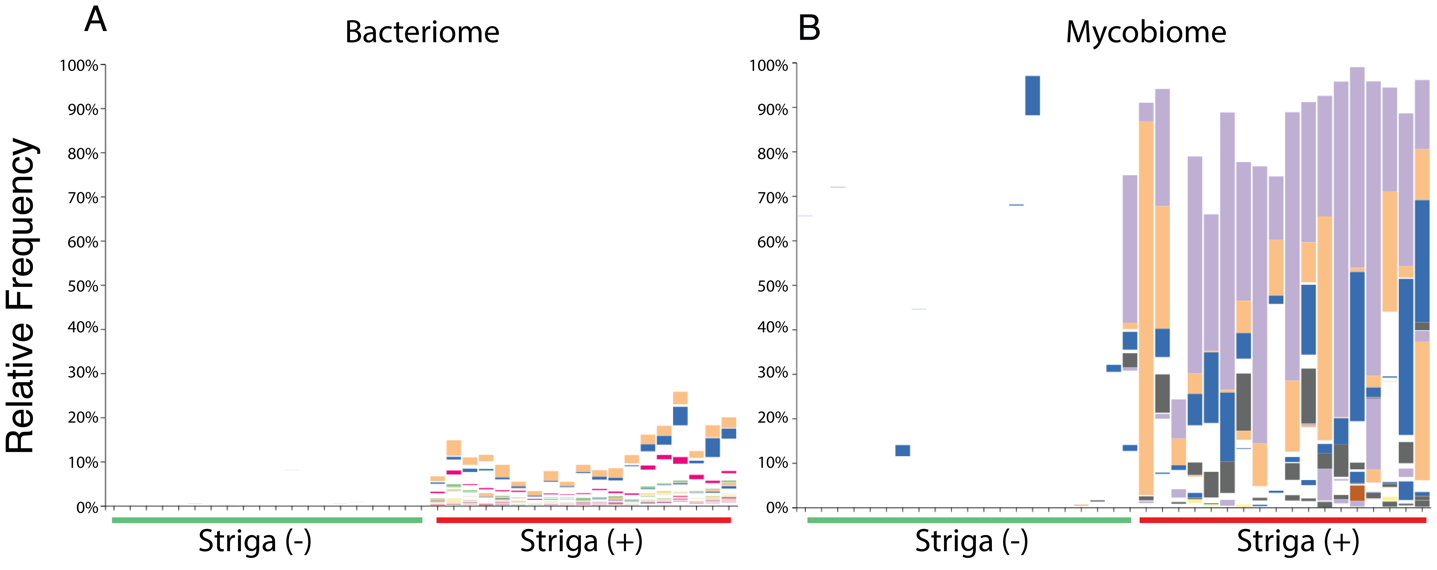

**Supplementary Figure SF7**. The relative increase in abundance of the members of the bacteriome (**A**) and mycobiome (**B**) of sorghum rhizosphere in response to *Striga* presence (*Striga* (+)). The relative abundance of the different taxa is shown in the Y-axis.
